# Machine learning reveals adaptive maternal responses to infant distress calls in wild chimpanzees

**DOI:** 10.1101/835827

**Authors:** Guillaume Dezecache, Klaus Zuberbühler, Marina Davila-Ross, Christoph D. Dahl

## Abstract

Distress calls are an acoustically variable group of vocalizations ubiquitous in mammals and other animals. Their presumed function is to recruit help, but it is uncertain whether this is mediated by listeners extracting the nature of the disturbance from calls. To address this, we used machine learning to analyse distress calls produced by wild infant chimpanzees. It enabled us to classify calls and examine them in relation to the external event triggering them and the distance to the intended receiver, the mother. In further steps, we tested whether the acoustic variants produced by infants predicted maternal responses. Our results demonstrated that, although infant chimpanzee distress calls were highly graded, they conveyed information about discrete events, which in turn guided maternal parenting decisions. We discuss these findings in light of one the most vexing problems in communication theory, the evolution of vocal flexibility in the human lineage.

## Introduction

Distress calls are an acoustically variable group of vocalizations found across mammals and other animal taxa (MacLean 1985; Lummaa et al. 1998; Zeifman 2001; Newman 2007; Bornstein et al. 2017). Their function is most likely to recruit help (Soltis 2004; Lingle and Riede 2014) but it is uncertain whether caregivers can base their reaction by inferring the nature of the disturbance from the acoustics of the calls alone, or whether they must also rely on context. The current view is that distress calls represent an acoustically graded system, with no clear acoustic boundaries (Zeskind et al. 1985; Porter et al. 1986; Soltis 2004; Lingle et al. 2012). However, the fact that a call system is graded does not automatically disqualify it from categorical perception (Fischer 1998). For humans, similarly, there has been an ongoing debate as to whether infant and adult (Raine et al. 2018) distress call variants convey discrete information about the event experienced by the callers (Wiesenfeld et al. 1981; Brennan and Kirkland 1982; Fuller 1991; Gilbert and Robb 1996; Soltis 2004).

In chimpanzees, distress calls are mainly produced by infants (Plooij 1984; Goodall 1986; Bard 2000). They typically are short tonal and low-pitched sounds, whose fundamental frequency and call rate may increase throughout the sequence (see SI for video examples; see SI Supplementary Figure 1 for spectrogram), potentially reflecting a difference in arousal and other internal states (Briefer 2012). In previous longitudinal research with free-ranging chimpanzees (Plooij 1984), distinctions have been made between (a) whimper-hoos (soft and low-pitched sounds rarely produced in sequences), (b) whimpers (sequences of noiseless sounds) and (c) ‘crying’ (which name originates for the fact resembles adult screams and is marked by rapid fluctuations in frequency). Apart from work on idiosyncratic distress calls (with features of the fundamental frequency contributing to individual distinctiveness) (Levréro and Mathevon 2013), no acoustic analyses have been conducted on chimpanzee distress calls. In this study, we used machine learning to revisit the hypothesis that context-specific acoustic variants in distress calling can vary with infant needs and predict maternal responses. Recent developments in machine learning have proven useful in the study of animal vocal communication (Mielke and Zuberbühler 2013; Fedurek et al. 2016; Turesson et al. 2016; Versteegh et al. 2016), but this technique has not yet been applied to distress calls and other graded animal signals given to discrete events (Barajas-Montiel and Reyes-García 2006; Saraswathy et al. 2012; Chang and Li 2016). We analysed calls from all accessible infants of an entire cohort of wild chimpanzees (*Pan troglodytes schweinfurthii*) (N=8) in the Sonso community of Budongo Forest, Uganda, during spontaneous interactions with their mothers (see SI Supplementary Table 1 for infants’ demographics). We extracted a large number of calls and subjected them to an automated feature extraction algorithm before training a supervised learning algorithm for subsequent categorisation. The procedure consisted in training a model to segregate calls given in a particular context. This model was then used to categorise new exemplars (Mohri et al. 2012), which enabled us to evaluate whether acoustically graded distress calls encode information about the context of emission. We also evaluated whether they encoded information about the distance between infant and mother, following reports in other primates (Bayart et al. 1990; Wiener et al. 1990). Finally, we looked at whether maternal responses can be predicted from the call structure alone, establishing a common acoustical code between chimpanzee infants and mothers.

**Figure 1.**
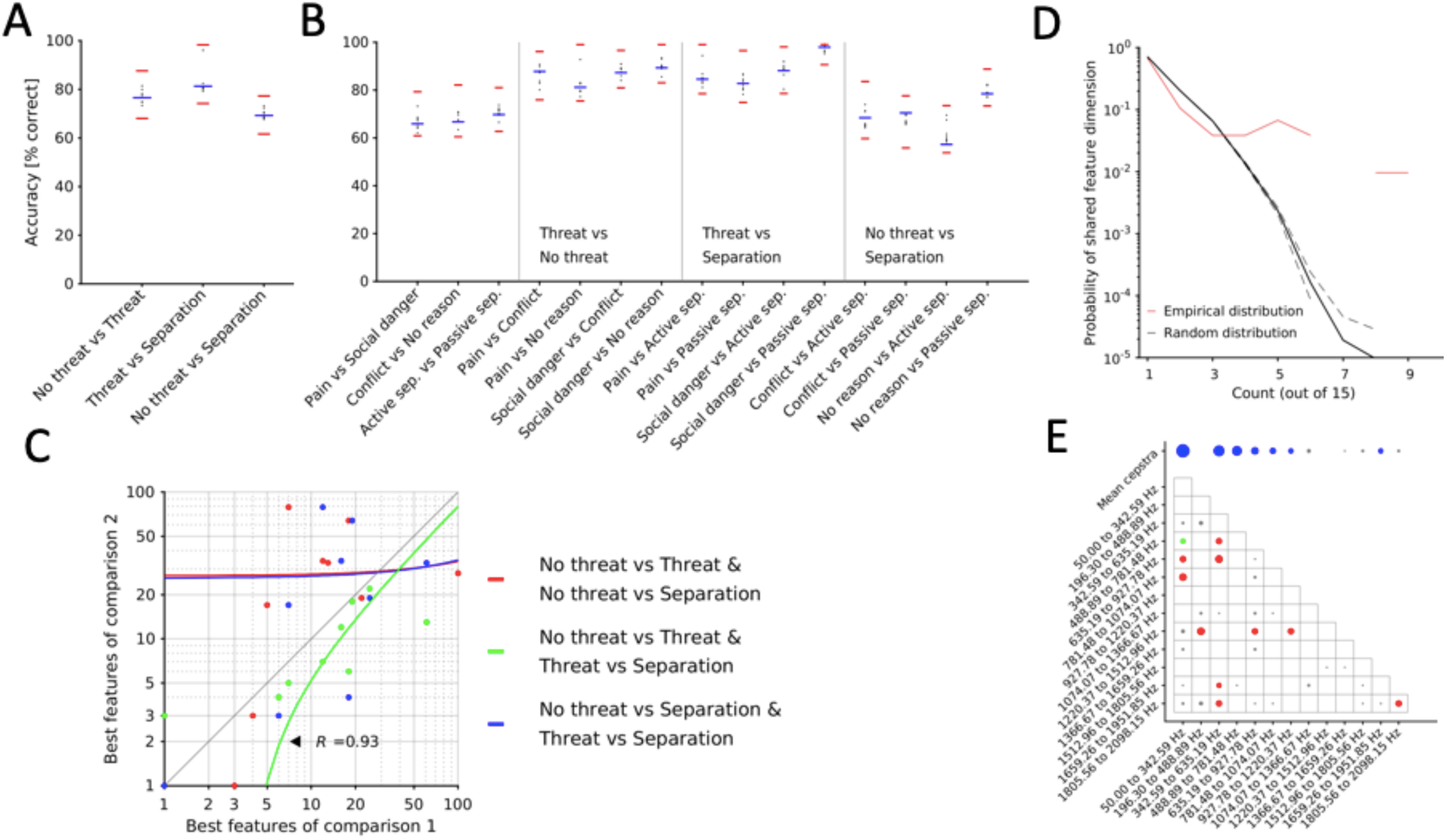
Model performance to classify across situations and infants’ problems and evaluation of key features contributing to classification (A, B) Accuracy values of classification runs are shown for all contrasts between situations (panel A) and problems (panel B). The blue horizontal bars indicate the means, the black dots the results from cross-validation and the red horizontal lines the minimal and maximal values of the leave-one-out procedure. (C) The feature set accounting best for individual contrasts are compared and correlated. Dots show individual samples; relationships between them are indicated by regression lines. (D) Feature evaluation procedure, showing the occurrence count (x-axis), reflecting the number of times a particular feature dimension was among the best 10 feature dimensions across all comparisons, and the relative frequency of n-counts (y-axis). In red, the empirical distribution is shown, in black a random distribution (solid line) and 95% confidence-interval in dotted lines. N-counts of 4 and more are significantly over-represented. (E) Significant feature dimensions are colour-coded and structurally aligned in a frequency-transformed representation. Blue dots indicate significant mean cepstra, red dots positive co-variances of cepstra, and green dots negative co-variances of cepstra. Grey dots indicate non-significant feature dimensions. The size of dots indicates the relative importance: the larger the dot the more frequently a feature dimensions has been used across all comparisons.

## Methods

### Subjects

Data were collected from infants (N=8) of the well-habituated wild chimpanzees (N≈70) of the Sonso community (Reynolds 2005) of Budongo Forest, Uganda, during February-June 2014, December 2014, March-June 2015 and April-May 2016. For further information about the study site, see (Eggeling 1947; Reynolds 2005). We received permission from the Ugandan Wildlife Authority (UWA) and the Uganda National Council for Science and Technology (UNCST) to conduct this study. SI Supplementary Table 1 presents a list of infants, their estimated birthdates, minimum and maximum ages in months over the course of the study, sex, number of contributed call episodes and number of extracted calls.

### Data collection

Distress calls were collected in all-occurrence fashion (Altmann 1974). Given their clear acoustic differences with other infant vocalizations (grunts, barks) and despite their acoustical variability, distress calls can be clearly identified by trained listeners due to their short, tonal, and sequential production pattern (see SI Supplementary Figure 1 for a spectrogram and SI for video and audio examples). Call episodes were video-recorded using a Panasonic HC X909/V700 video-camera, with the sound captured by a Sennheiser MKE-400 shotgun microphone. To account for any missing information in the videos, AUTHOR1 recorded all potential causes of infant calls, the reactions of the mother, and the distance between them. In our study, the infants’ perceived needs were classified according to their situation and problems (see SI Supplementary Table 2). Distance between mother and infant at the onset of the call was also determined and was coded as ‘Supported’ if most of the weight of the infant was then supported by the mother; ‘Contact’ if there was physical contact with mother but without full support (e.g., standing on the ground); ‘Armsreach’ if the infant was within the arm’s reach of the mother, and ‘Beyond’ if the infant was beyond the arm’s reach of the mother. AUTHOR1 also commented on the reaction of the mother from the onset of the call and up to 10 seconds after the call offset or when the infant was not visible anymore; AUTHOR1 further reported whether the mother gazed towards the infant (as evidenced from the facial orientation), approached the infant, collected the infant or vocalized in response to the infant’s distress calls. Distance between the mother and her infant at call onset and mothers’ reaction were intra-reliably validated on 19.5% of all call episodes (κ distance = 1; κ gaze = 0.90; κ approach = 0.87; κ collection = 0.96; κ vocalization = 0.92). Since the situations and problems attached to infant calling were heavily dependent on external events not necessarily visible in the videos, we mostly relied on live commentaries and annotations by AUTHOR1 to code them.

### Quantification and statistical analysis

Acoustic analysis was performed with Matlab (Mathworks Inc., Natick, MA, USA) and contained the following steps: (1) call extraction, (2) features extraction, (3) feature selection, (4) call classification, and (5) feature evaluation.

#### Call extraction

We first pre-processed the raw audio files by applying a band pass filter from 100 to 700 Hz and filtering out background noise using a Vuvuzela denoising algorithm (Boll 1979; Ephraim and Malah 1984). Distress calls can grade beyond a frequency of 750 Hz but this was the maximal value of the peak fundamental frequency at mid-call of the first distress call of the series we collected. This pass band was, therefore, appropriate for the localization of calls and to avoid false positives. We used a voice-activity detection (VAD) algorithm to read out vocalization on- and offsets. We therefore subdivided the audio files into 10ms segments, determined the prominence of energy of each call time element and applied a cut-off threshold at the 80^th^ percentile of the energy-prominence distribution. We further calculated the spectral roll-off points, defined as the power spectral distribution, with a threshold at the 85^th^ percentile. From the remaining extractions, we then sorted out those extractions that were shorter than 40ms or longer than 120ms (corresponding to the range of durations extracted manually from whimper-like calls, based on the first call of the distress call series we collected) and subjected the remaining extractions to a human-based validation process, leaving 1330 call on- and offsets. We extracted the full frequency spectrum ranging from 50 to 4000 Hz at given call on- and offsets.

#### Feature extraction

Feature extraction is the extraction of a subset of relevant features from each call to minimize data size, redundancy in information and computational efforts and increase generalization ability of a classifier (Tajiri et al. 2010). Similar to (Mielke and Zuberbühler 2013; Fedurek et al. 2016), we extracted mel frequency cepstral coefficients (MFCCs), representing the envelope of the short-time power spectrum as determined by the shape of the vocal tract. We subdivided the vocalizations into windows of 25ms segments and 10ms steps between two successive segments, to account for signal changes in time. We warped 26 spectral bands and returned 13 cepstra, resulting in feature dimensions of 13 values each. We then calculated the mean and co-variances of each cepstra over the collection of feature segments, resulting in a 13-value vector and a 13 x 13-value matrix, respectively, and concatenated to 104-unit vectors. We also applied feature scaling to values between 0 and 1.

Prior to classification, we conducted a feature selection procedure, reducing the number of features to a set of reliable features, explaining most of the variance in a given data set. We applied a *t*-test on each feature dimension comparing values of a given feature dimension sorted by predefined classes labels (e.g. One type of situation vs. another). For each comparison we used randomly determined 75% of the samples and re-ran the procedure five times for each comparison. Such feature selection procedure is called a filter approach, where general characteristics are evaluated for the selection without subjecting the dimensions to a classifier.

#### Call classification

For the classification of calls, we implemented support vector machines (SVMs) using the LIBSVM toolbox (Chang and Lin 2011). Classification routines consisted of training and testing phases: First, separate training samples (i.e. feature vectors of extracted calls) were labelled with an attribute (e.g. Situation A = 0, Situation B = 1) and used to build a model of support vectors that optimally separate the two classes. Testing samples without attribute labels were then used to measure the model’s performance to generalize for novel samples. We used a radial basis function (RBF) kernel and 5-fold cross-validated the parameters C and gamma with separate smaller data sets (see (Fedurek et al. 2016) for details on the procedure). Training was performed on 80% of the data samples, testing with the remaining 20%. We used the 40 best feature dimensions for classification, omitting all others. Training and testing were always done on two classes. We obtained performance scores from models trained and tested on the same labels (basic classification procedure) and from cross-comparisons of conditions, such as training on situations and testing on one type of maternal reaction. This latter way of system evaluation has been previously described and provides useful insights into the nature of information coding (Caldara and Abdi 2006; Dahl et al. 2016). Generally, the goal was not to find the classifier achieving the highest performance scores from a machine learning point-of-view, rather than evaluating relative performance differences and the relationship of features contributing to one task as opposed to another.

#### Feature evaluation

For call situations, we determined the extent to which comparisons (no threat vs. threat; no threat vs. separation; threat vs. separation) share similar feature dimensions. We determined the 10 best feature dimensions of each comparison, as outlined above, and correlated the feature numbers of those feature dimensions in pairwise fashion, using Spearman’s rank correlation. Feature numbers refer to the topological organisation of the Mel-frequency space; hence close feature numbers are indicative for similar structural features accounting for both comparisons. To determine the feature dimensions that are particularly critical for the classification of distress calls, we assessed whether feature dimensions have been repeatedly used by the classifier overall in the classification of distress calls. We therefore considered the 15 types of comparisons regarding infants’ problems (e.g., “Conflict vs No reason”). Regarding the distance between infant and mother, we considered six types of comparisons (e.g., “Supported vs Contact”). Regarding the mother’s response, we considered four yes-no comparisons, namely for “Gaze”, “Approach”, “Collection” and “Vocalization”. We then determined the empirical distribution of the ten features achieving best classification power out of the 40 feature dimensions that resulted from the feature selection algorithms (see above). The total of 10 best features was an arbitrary choice. As a baseline, a random distribution of “best features” for each comparison was determined by randomly selecting 10 out of 104 features. The frequency distribution across all comparisons were determined and 95% confidence intervals were calculated by running the procedure for 1,000 times. We then reconstructed the underlying frequency bands of significant feature dimensions, resulting in “feature maps” for “situation”, “distance [between infant and mother]” and “maternal response”. In a further step, we calculated differences in feature importance between “situation”, “distance [between infant and mother]” and “maternal response” by pairwise contrasting feature maps. We contrasted two feature maps by subtracting corresponding frequency values, reflecting the occurrence of one particular mean cepstra or co-variance of two cepstra, if at least one of the corresponding two values from the two feature maps was significant. The reason for this is that subtracting two non-significant values would be meaningless.

### Analysis

We obtained performance scores in the form of correct classification. We also obtained a performance baseline, which was determined by always predicting the most frequent label in the training set. We then contrasted the classifiers performance scores with those of the baseline using *t*-test. Additionally, to ensure that no single individual determined the outcome of the classification routine, a leave-one-out method was used where the general classification procedure was re-run eight times, omitting calls produced by one individual in each run. We tested for a significant interaction between leave-one-out runs and the individual comparisons using a two-way ANOVA.

## Results

### Distress calls encode information about the situation and infants’ problems

Infants emitted distress calls when separated from their mother (situation: separation), when exposed to a danger such as aggression or the experience of pain (situation: threat) or for reasons not related to separation nor threat (situation: no threat). Classification accuracy for all 3 situations was high and significantly higher than baseline (no threat vs. threat: M_emp_ = 74.50, SD_emp_ = 4.26; M_base_ = 59.61, SD_base_ = 4.56; *t*(18) = 11.47, *p* < 0.001; threat vs. separation: M_emp_ = 81.40, SD_emp_ = 4.03; M_base_ = 58.19, SD_base_ = 4.39; *t*(18) = 19.94, *p* < 0.001; no threat vs separation: M_emp_ = 71.02, SD_emp_ = 3.09; M_base_ = 58.48, SD_base_ = 3.82; *t*(18) = 9.85, *p* < 0.001; Figure 1A).

The results of a leave-one-out procedure (in which one of the infants was at turn removed from one of the 8 individual runs) was not consistent with effects due to single subjects (No significant interaction between situation and leave-one out runs: *F*(14,720) = 1.36, MS = 17.16, *p* = 0.17).

Can distress calls acoustics code for the exact problem the infants were dealing with? In some of the separation situations, physical distance could have been initiated by the mother and the infant started calling in response to this movement from the mother (problem: active separation). In other separation situations, the infant was already away from the mother and started calling in the absence of specific travelling movements from the mother (problem: passive separation).

Similarly, threat situations involved different problems. In some threat situations, infants seemingly experienced physical pain (e.g., when mother had been moving abruptly) (problem: pain). In others, a social or natural phenomenon appeared threatening to the infant (e.g., aggression in the vicinity) (problem: social danger).

Finally, no threat situations could expose infant to conflict problems (problem: conflict) such as when mother and infant disagree on travel decisions or on food provisioning (e.g., mother refusing infants’ access to the nipple). In other instances, no obvious trigger induced distress calls in the infant (problem: no reason).

Classification accuracy for problems turned out significantly higher than the baseline (all *p*-values < .05, see SI Supplementary Table 3) (Figure 1B). Interestingly, discrimination for problems that belonged to the same situation (e.g., problems ‘pain’ and ‘social danger’ are both part of situation ‘threat’) turned out to be reduced in accuracy, albeit being significantly different from the baseline (SI Supplementary Table 3). By contrast, the highest discrimination performances were found in contrasts where one of the detailed problems occurred in the threat situation and the other into the no threat situation. Similarly, contrasts where one of the problems fell into the threat situation and the other into the separation situation yielded high performance scores. Classification of contrasts involving problems across separation and no threat situations was more modest, albeit significantly higher than baseline level (SI Supplementary Table 3). Those contrasts score significantly lower than those comparing problems across threat and other situations (threat vs. no threat: M_1_ = 85.30, SD_1_ = 6.14; M_2_ = 68.46, SD_2_ = 8.25; *t*(78) = 14.13, *p* < 0.001; threat vs separation: M_1_ = 87.74, SD_1_ = 8.39; M_2_= 68.46, SD_2_ = 8.25; *t*(78) = 16.19, *p* < 0.001).

### How distress calls encode information about the situation

Feature analysis revealed that the classification between pairs of situations involving threat relied on similar feature dimensions in similar fashion, indicated by a similar ranking of those features for contrasts threat vs. no threat and threat vs. separation. The correlation of feature numbers at given rank in both contrasts yielded a positive correlation (*r*_*s*_ = .93, n = 10, *p* < .001, Figure 1C). This was not the case for pairs of contrasts comparing separation and other situations, and those comparing no threat and the two other situations (*r*_*s*_ = .47 and *r*_*s*_ = .61 respectively; Figure 1C).

We further examined the feature dimensions that accounted most for the classification of distress calls according to the specific problems the infants were exposed to. We determined the point at which an empirical number count, i.e. the number of times a particular mean cepstra or a covariance of two frequency bands contributed significantly to the classification of distress calls, significantly surpassed a random distribution of number counts, relative to the total number of comparison (here 15) (Figure 1D). In other words, features that are of significance for four and more comparisons are significantly different from the baseline. Those feature dimensions are marked in Figure 1E, and emphasize the importance of the first, third and fourth mean cepstra, accompanied by covariances between frequency bands around 342.59 to 1074.07 Hz and 50 to 781.48 Hz.

### Distress calls encode information on the distance between infant and mother at call onset

At call onset, the infants could either be completely supported (distance: supported) or in contact (distance: contact) with the mother. They could also be further away from their mother, within or beyond her arms’ reach (distance: armsreach; distance: beyond, respectively). Based on distress calls acoustics, our model classified distance between infant and mother at levels higher than baseline (supported vs. contact: M_emp_ = 64.16, SD_emp_ = 2.52; M_base_ = 56.85, SD_base_= 2.99; t(18) = 5.16, p < 0.001; supported vs. armsreach: M_emp_ = 85.70, SD_emp_ = 2.43; M_base_ = 58.05, SD_base_ = 5.45; t(18) = 24.37, p < 0.001; supported vs. beyond: M_emp_ = 96.13, SD_emp_ = 3.24; M_base_ = 56.49, SD_base_ = 4.40; t(18) = 36.65, p < 0.001; contact vs. armsreach: M_emp_ = 88.95, SD_emp_ = 3.05; M_base_ = 57.00, SD_base_ = 4.34; t(18) = 29.04, p < 0.001; contact vs. beyond: M_emp_ = 95.91, SD_emp_ = 4.03; M_base_ = 55.15, SD_base_ = 3.97; t(18) = 37.65, p < 0.001; armsreach vs. beyond: M_emp_ = 79.84, SD_emp_ = 3.32; M_base_ = 59.23, SD_base_ = 2.63; t(18) = 18.29, p < 0.001; Figure 2A). This was not due to single subjects (No interaction between distance and leave-one-out runs: *F*(5,288) = 0.92, MS = 17.44, *p* = 0.47).

**Figure 2.**
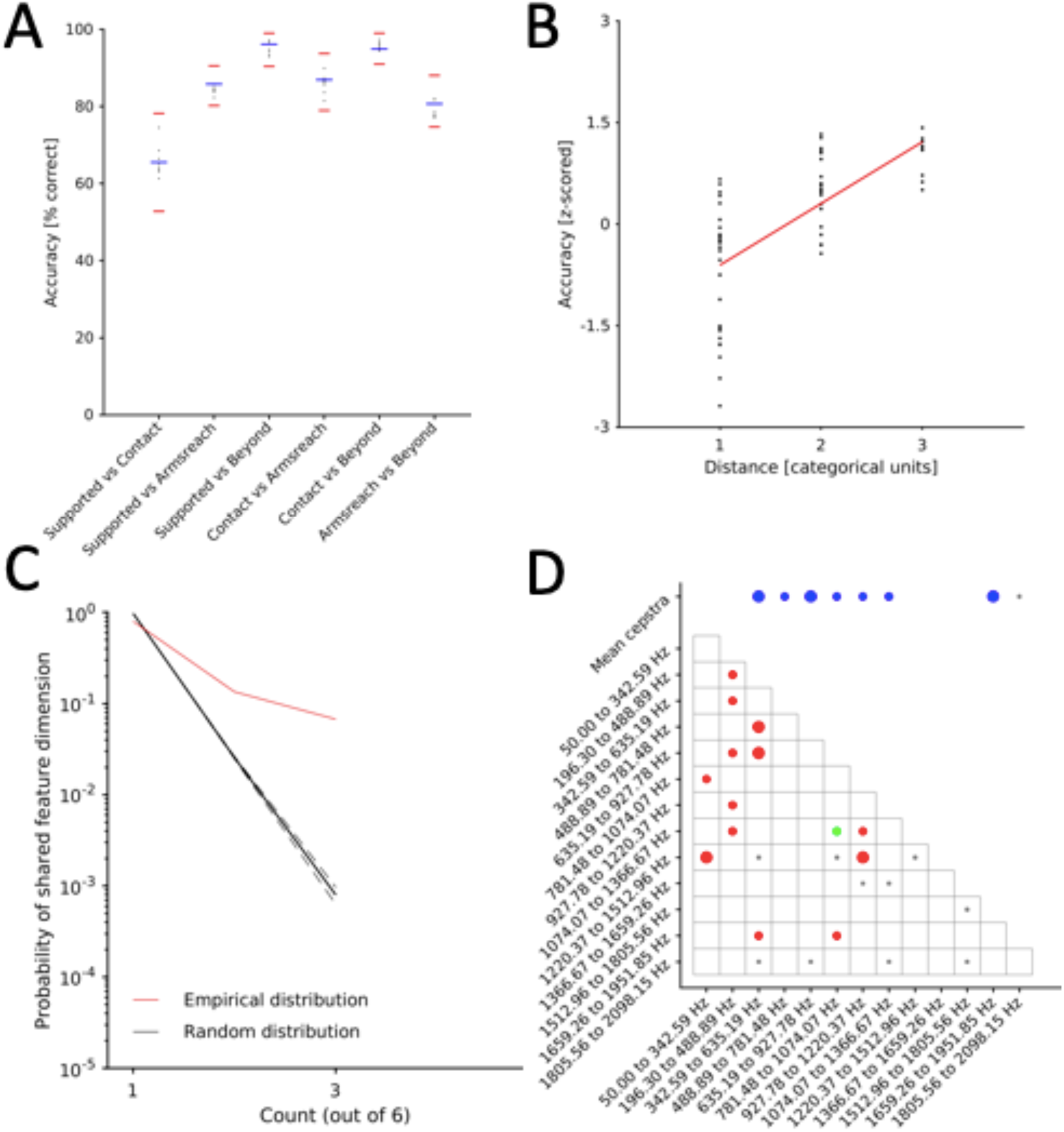
Model performance to classify across distance between mother and infants and evaluation of key contributing features. (A) Accuracy values of classification runs are shown, contrasting various categorical distances between infant and mother. Colour-code as in panels Figure 1A&B. (B) Accuracy scores are plotted according to the relative distance of distance labels compared. Accuracy values are shown as z-scores. Red line represents the linear regression fit. (C) Feature evaluation procedure, showing the occurrence count (x-axis), reflecting the number of times a particular feature dimension was among the best 10 feature dimensions across all comparisons, and the relative frequency of n-counts (y-axis). Colour-code as in Figure 1D. N-counts of 2 and more are significantly over-represented. (D) Significant feature dimensions (see Figure 1E for colour codes).

We further contrasted distance classes, assigning categorical units of distance, with 1 being the most proximal (supported) and 4 being the most distal (beyond). We then binned classification scores according to the relative distance of the classes compared (e.g., supported vs. contact was binned with ‘contact vs. armsreach’ (relative distance of 1)) and z-scored the binned values. We found a significant increase of classification accuracy as relative distance between the two distances compared increased (F(2,59) = 35.12, MS = 16.21, p < .001; Figure 2B). This suggests that variation in classification performance (range: 64-96 % correct classification) reflects a systematic tendency to code for distance in the structure of infant chimpanzee distress calls, where calls emitted at relatively similar distance to the mother are harder to acoustically discriminate than those emitted at larger relative distance to each other.

Examination of features dimensions that most contribute to classification revealed that features that are of significance for two and more comparisons are significantly different from the baseline. Indeed, the empirical number count 2 significantly surpassed a random distribution of number counts, relative to the total number of comparison (here 6) (Figure 2C). The third and fifth mean cepstra, accompanied by covariances between frequency bands in the lower to mid-range of the Hz-spectrum, most contribute to classification (Figure 2D).

### Distress calls acoustics predicts maternal intervention

If calls contain information about the nature of the external event triggering infants’ distress and about the distance to their mothers, can they predict maternal response? Four types of maternal responses were examined for their occurrence in response to infant distress calling: gaze towards the infant (maternal response: gaze), approach (maternal response: approach), collection of the infant (maternal response: collection) or vocal response (maternal response: vocalization).

The model discriminated accurately and above baseline the probability of the mother to gaze towards (M_emp_ = 95.13, SD_emp_ = 4.48; M_base_ = 58.09, SD_base_ = 4.14; *t*(18) = 33.70, *p* < 0.001), approach (M_emp_ = 85.79, SD_emp_ = 3.13; M_base_ = 57.79, SD_base_ = 4.18; *t*(18) = 25.14, *p* < 0.001) collect the infant (M_emp_ = 62.58, SD_emp_ = 2.91; M_base_ = 60.03, SD_base_ = 3.15; *t*(18) = 0.20, *p* < 0.05) and to vocalize in response to the distress calls (M_emp_ = 73.83, SD_emp_ = 2.13; M_base_ = 55.39, SD_base_ = 2.24; *t*(18) = 16.75, *p* < 0.001) (Figure 3A). Although discrimination was always above baseline level, it was more modest to predict maternal collection and vocalization behaviours. The relatively low number of observations of vocal response from the mother can explain this latter result (See SI Supplementary Figure 2A). A leave-one-out procedure revealed no significant interaction between individual runs (N = 8) and maternal response comparisons (No significant interaction between maternal response and leave-one out runs: *F*(3,192) = 0.67, MS = 1.87, *p* = 0.57).

**Figure 3.**
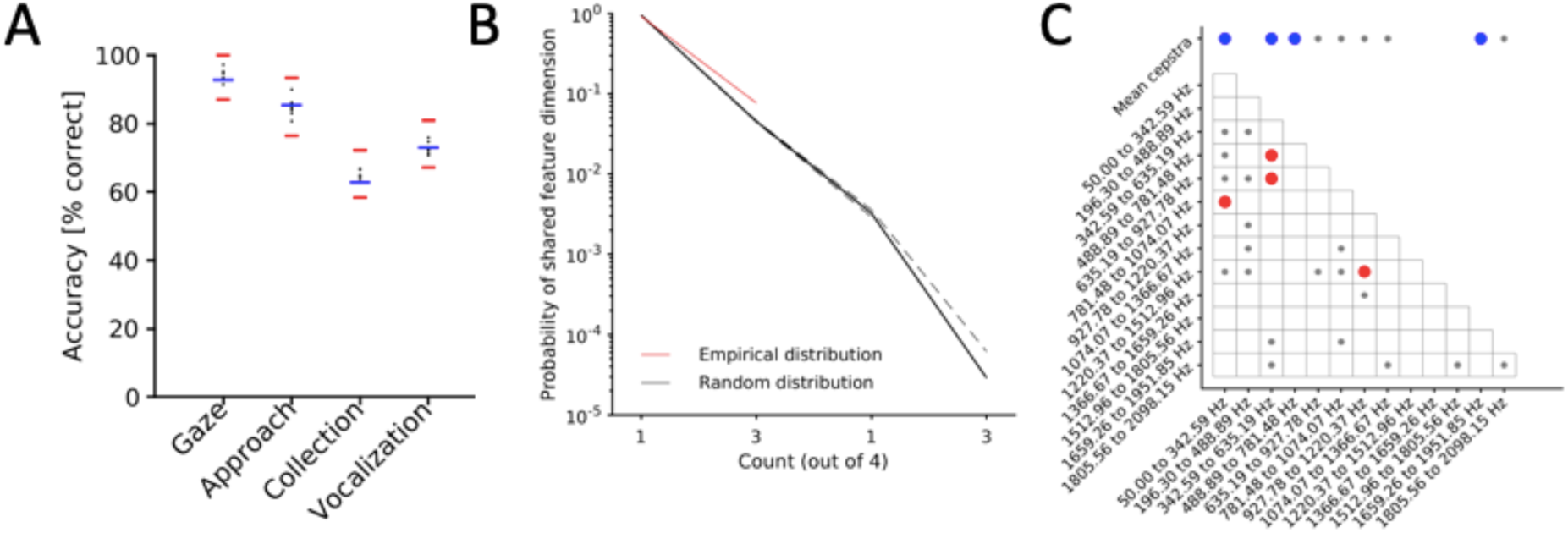
Classification performance per type of maternal intervention and evaluation of contributing features. (A) Accuracy values of classification runs for maternal intervention. Colour-code as in Figures 1A&B and 2A. (B) Feature comparison, showing the occurrence count (x-axis), reflecting the number of times a particular feature dimension was among the best 10 feature dimensions across all comparisons, and the relative frequency of n-counts (y-axis). Colour-code as in Figure 1D and 2C. N-counts of 2 and more are significantly over-represented. (C) Significant feature dimensions are colour-coded and structurally aligned in a frequency-transformed representation, following Figure 1E and 2D.

To evaluate the features most contributing to determining whether mother did or did not react to infant calls, we plotted the mean expression of each feature dimensions for both the presence (Yes) and the absence (No) of a maternal behaviour. We found that selected features were often located at the outer edge of the distribution (See SI Supplementary Figure 2B), i.e., they account for one class significantly stronger than for the other class. The contrasts that yielded high performance scores showed more distinct feature separation (e.g., Gaze plot of SI Supplementary Figure 2B) than those contrasts that yielded rather modest classification (e.g. Collection plot of SI Supplementary Figure 2B). We also examined the feature dimensions that accounted most for the classification of distress calls according to maternal response category labels and found that features that are of significance for two and more comparisons are significantly different from the baseline. The empirical number count 2 significantly surpassed a random distribution of number counts, relative to the total number of comparisons (here 4) (Figure 3B). The importance of the first, third, fourth as well as the 12th mean cepstra, accompanied by covariances between frequency bands around 342.59 to 1074.07 Hz and 50 to 781.48 Hz (Figure 3C). The feature distribution of maternal response classification, therefore, resembles the feature distribution of problems faced by the infants (Figure 1D).

### Mother-infant interactions

The number of approaches and collections was higher in the most critical situation (i.e., threat) (See SI Supplementary Figure 2A), when problems were social danger and pain. To further understand how information-coding of the situation in the distress calls contribute to trigger adaptive maternal response, we trained a model on discriminating situations and infants’ problems and tested its performance in classifying the presence of the various types of maternal responses. Our model discriminated significantly higher than baseline between threat and no threat situations across all maternal response types (M_emp_ = 69.38, SD_emp_ = 9.39; M_base_ = 54.65, SD_base_ = 2.87; *t*(78) = 9.48, *p* < 0.001), as well as between threat and separation situations (M_emp_= 72.31, SD_emp_ = 8.87; M_base_ = 55.33, SD_base_ = 2.67; *t*(78) = 11.57, *p* < 0.001) (Figure 4A). However, contrasts based on the two other situations (no threat and separation) were not successfully discriminated based on the maternal responses (M_emp_ = 55.33, SD_emp_ = 4.33; M_base_= 54.90, SD_base_ = 2.87; *t*(78) = 0.52, *p* = 0.300) (Figure 2D). Testing the model with specific problems infant were exposed to, we found all comparisons to be significant (*p* < .05, SI Supplementary Table 4) but two (contrasts no reason vs. active separation, and conflict vs. no reason (Figure 4B). Gaze was a relatively reliable action of the mother, particularly during problems involving a threat situation. On the other hand, collection and vocalization were found to be more modest in discriminating between infants’ problems.

**Figure 4.**
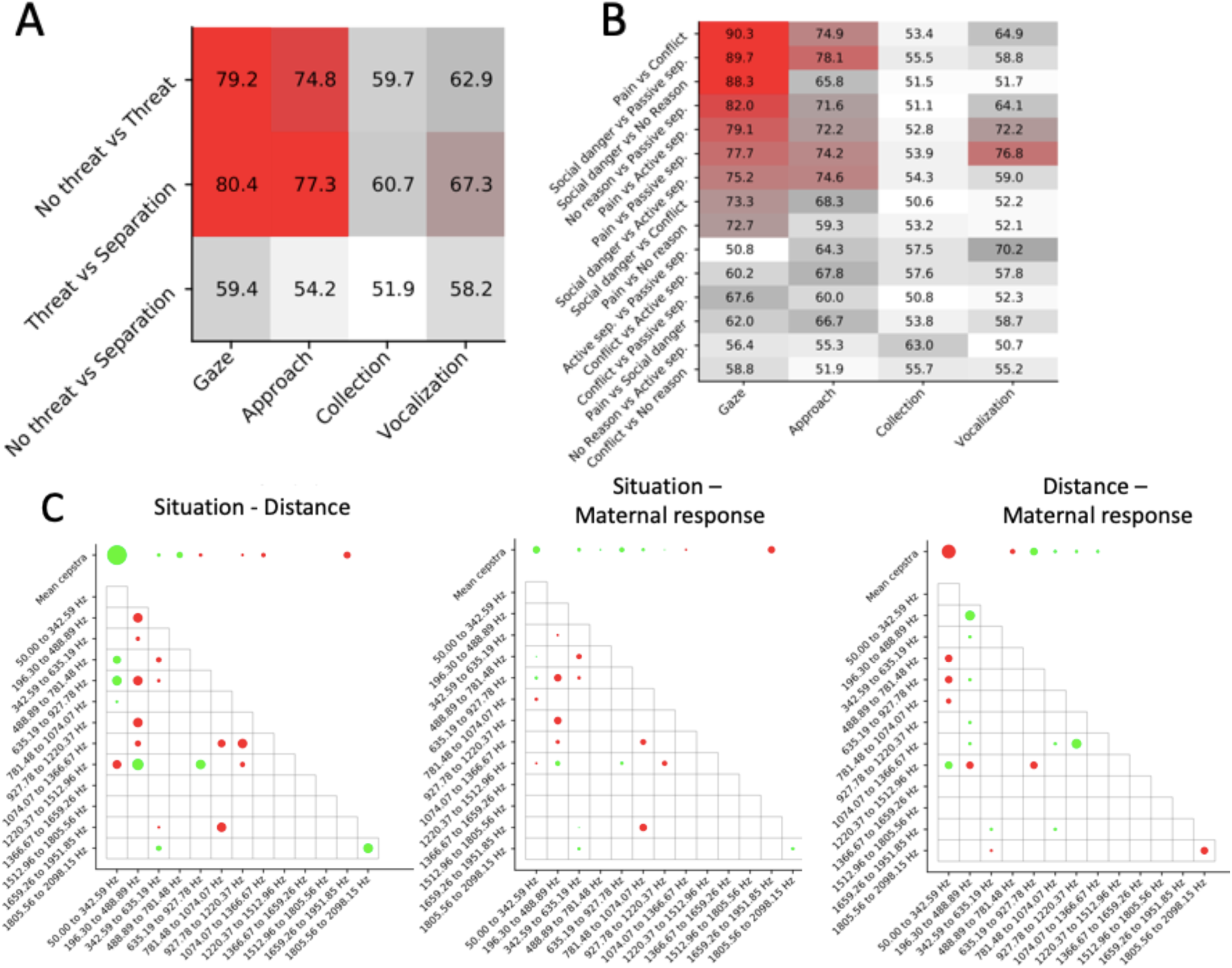
Mother-infant interaction classification models. (A & B) Accuracy values of classification runs for models trained on situation (A) or infants’ problems (B) contrasts (y-axis) and tested on maternal intervention contrasts (x-axis). The colour represents the level of accuracy (0% corresponds to the white colour; 100% corresponds to the red colour). (C) Feature map comparison. Feature maps as shown in Figure 1E, 2D and 3C were contrasted by subtracting corresponding frequency values, reflecting the occurrence of one particular mean cepstra or co-variance of two cepstra, if at least one of the corresponding two values from the two feature maps was significant. Green indicates greater importance for the first term (e.g., ‘situation’ in the extreme-left panel), red for the second term of the comparison (e.g., ‘maternal response’ in the extreme-right panel).

To address the extent to which the structural differences of distress calls produced by the infants across situations reflect structural differences encoding distance and predicting maternal response, we pairwise contrasted feature maps for situation, distance and maternal response. We found that, when comparing situation and distance, certain features were selective for situation that related strongly to the first frequency band ranging from 50 to 342.59 Hz and co-varied with frequency bands ranging from 488.89 to 927.78 Hz. Also, the frequency band centring at 1366.70 Hz co-varied with the second frequency band ranging from 196.30 to 488.89 Hz and the fifth ranging from 635.19 to 927.78 Hz (Figure 4C). On the other hand, distance was selectively coded in the second frequency band ranging from 196.30 to 488.89 Hz, co-varying with multiple feature dimensions (Figure 4C). In the comparison between situation and maternal response (Figure 4C), we found similar components involved, however, to a marginal degree, indicating that there are similar main features accounting for most of the classification of distress calls for both the situation and maternal response. Comparison between distance and maternal reflected the comparison between distance and situation (Figure 4C).

## Discussion

In this research, we tackled a long-standing, widely-debated and cross-disciplinary question, i.e., whether highly graded infant distress cries carry different meanings that guide parenting decisions.

Distress calls have been described as acoustically continuous or graded and various studies (notably in humans) have raised the possibility that these calls convey discrete information (Müller et al. 1974; Wiesenfeld et al. 1981; Brennan and Kirkland 1982; Fuller 1991; Soltis 2004). Here, we used supervised machine learning to investigate distress calling behaviour in chimpanzees, our closest extant relative together with bonobos, to attempt to answer whether infant distress calls contain information about the nature of the external events experienced by the caller, its context (i.e., distance with the mother) and whether they drive maternal interventions.

We found that distress calls contain information about the type of external events triggering calling as well as the nature of the problem. In particular, we found that a model trained on discriminating a threatening situation from others was better at predicting the subsequent response of the mother, suggesting that distress calling during threat situations rely on specific acoustic features. This is further exemplified by the fact that discriminations between distress calling produced in threat situation vs. separation situation on one hand, and threat vs. no threat situations on the other hand, relied on common features.

We also found that the distance between infants and their mothers at distress call onset modulate distress call structure, with calls being acoustically most distinctive as relative distance between infants and mothers increases.

Another finding was that a mother’s reaction could be predicted based only on the acoustic characteristics of her infant’s calls, that is, when she would look towards the infant, approach it, collect it, or simply respond by vocalizing. Further, we found that feature maps supporting the classifications of calling situations and maternal response are relatively similar, i.e., that the most critical feature dimensions in call production also contribute to call interpretation by the mother. Together, this suggests that, in principle, chimpanzee mothers can rely exclusively on the acoustic information provided by their infants, to make intervention decisions based on the degree of urgency of the infants’ request. This ability must be particularly important in a species where infanticide by both males and females is commonly reported (Arcadi and Wrangham 1999; Newton-Fisher 1999; Watts and Mitani 2000; Townsend et al. 2007; Lowe et al. 2019*a*, 2019*b*), potentially enhancing the fitness of individuals able to convey specific needs in their vocalizations.

We interpret our findings as evidence for a common code between chimpanzee infants and their mothers, helping mothers to adjust their behavioural responses to the specific needs of their infants. Nevertheless, it is clear that infant distress calls are often given together with other signals (notably gestural and postural signals (Hobaiter and Byrne 2011, 2014; Fröhlich et al. 2016, 2018; Fröhlich and Hobaiter 2018)) and are interpreted by recipients in terms of the perceived dangerousness of the situation. How recipients consider and integrate across these different sources of information needs to be addressed by future research. This being said, our study shows that distress calls in chimpanzees are acoustically rich enough to convey information about the external events triggering the calls, similar to classic studies on primate alarm calls, and as such help caregivers to shape evolutionarily important intervention decisions.

## Data availability

Data are available upon reasonable request to the last Author.

## Acknowledgments

We thank the Ugandan Widlife Authority (UWA) and the Ugandan National Council for Science and Technology (UNCST) for permission to conduct the study, Geoffrey Muhanguzi, Caroline Asiimwe, Sam Adue and Monday M’Botella for their support in the field. We are grateful to the Royal Zoological Society of Scotland for providing core funding to the Budongo Conservation Field Station.

This research was supported by a Fyssen fellowship awarded to GD, funding from the European Union’s Seventh Framework Programme for research, technological development and demonstration (grant agreement no 283871), and the Swiss National Science Foundation (PZ00P3_154741) awarded to CDD.

## Supplementary Information (SI)

### Video and acoustic examples of distress calls

https://youtu.be/vRV-Fme_pjY

Foreground: KO moves away from the mother (KL). He then emits distress calls (around 8s) and reunites with KL.

Background: KC (KL’s oldest son) plays with OZ. By 11s, OZ emits distress calls after rough play with KC. OZ’s mother (OK) intervenes.

https://youtu.be/yagdANhXF1Y

KJ producing distress calls after separation with his mother.

**Supplementary Table 1.**
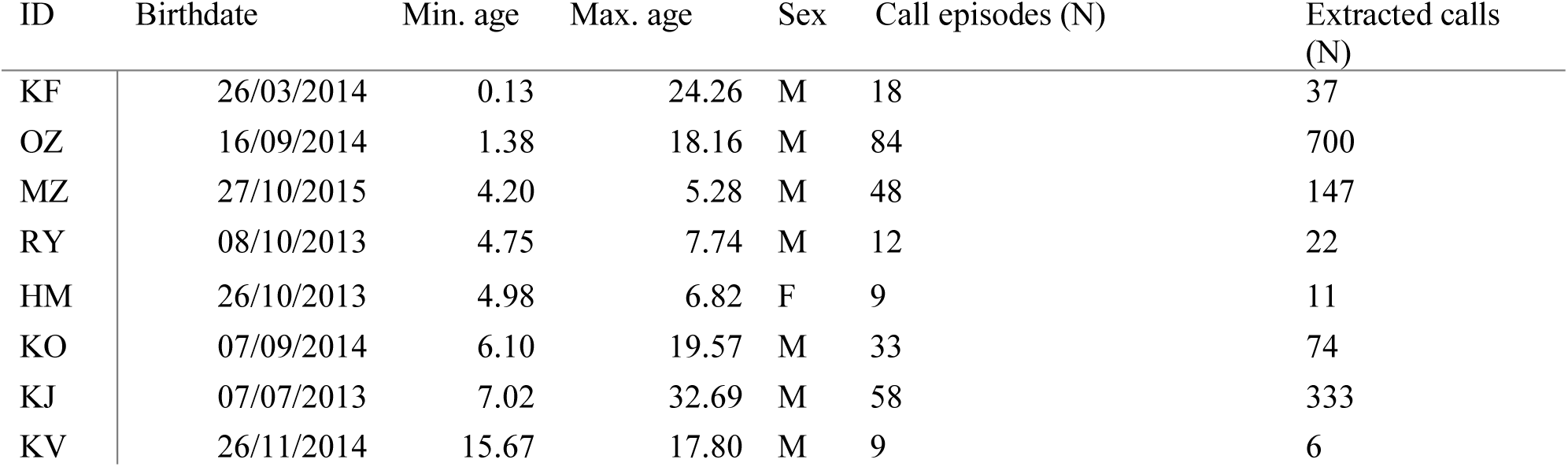
List of infants, estimated birthdate, minimum and maximum age in months over the course of the study, sex, number of contributed call episodes and number of extracted calls.

**Supplementary Table 2.** Situation and problems faced by the infant and definitions.

| Situation | Problem | Definition |
| --- | --- | --- |
| Threat | Social danger | When a natural phenomenon (suspicion of a dangerous animal in the surrounding or vegetation falling), a human observer or another chimpanzee appears threatening to the infant |
|  | Pain | When the infant experiences clear physical pain |
| Separation | Active separation | When the mother is initiating travelling without the infant |
|  | Passive separation | When mother and infant are already separated by 2 meters or more |
| No threat | Conflict | When mother and infant are engaged in a conflict of interest regarding travel decisions or food provision (begging for food or access to nipple) |
|  | No reason | When there is no detectable trigger that induces distress calls in the infant or mild discomfort |

**Supplementary Table 3.** Classification performances for the contrasts between problems faced by the infants, with mean (M) and standard deviation (SD) for both empirical and baseline distributions, and associated *t* statistics.

|  | Empirical | Baseline | Statistics |
| --- | --- | --- | --- |
| Pain vs Social danger | M = 63.30, SD = 4.77 | M = 57.03, SD = 4.57 | $t(18) = 2.65, p < 0.01$ |
| Conflict vs No reason | M = 67.46, SD = 2.54 | M = 58.53, SD = 4.06 | $t(18) = 6.31, p < 0.001$ |
| Active separation vs Passive separation | M = 70.51, SD = 3.92 | M = 58.70, SD = 1.21 | $t(18) = 9.56, p < 0.001$ |
| Pain vs Conflict | M = 85.29, SD = 4.84 | M = 57.10, SD = 5.05 | $t(18) = 24.35, p < 0.001$ |
| Pain vs No reason | M = 80.05, SD = 4.53 | M = 57.05, SD = 4.83 | $t(18) = 19.37, p < 0.001$ |
| Social danger vs Conflict | M = 88.33, SD = 5.10 | M = 57.65, SD = 2.86 | $t(18) = 27.47, p < 0.001$ |
| Social danger vs No reason | M = 90.48, SD = 3.90 | M = 55.28, SD = 3.40 | $t(18) = 32.36, p < 0.001$ |
| Pain vs Active separation | M = 85.68, SD = 2.30 | M = 58.99, SD = 3.99 | $t(18) = 24.17, p < 0.001$ |
| Pain vs Passive separation | M = 81.26, SD = 4.74 | M = 58.91, SD = 3.94 | $t(18) = 18.97, p < 0.001$ |
| Social danger vs Active separation | M = 86.93, SD = 4.34 | M = 58.17, SD = 3.28 | $t(18) = 25.77, p < 0.001$ |
| Social danger vs Passive separation | M = 99.45, SD = 5.23 | M = 56.49, SD = 3.18 | $t(18) = 39.60, p < 0.001$ |
| Conflict vs Active separation | M = 65.98, SD = 4.33 | M = 57.92, SD = 4.71 | $t(18) = 4.55, p < 0.001$ |
| Conflict vs Passive separation | M = 69.52, SD = 5.43 | M = 59.82, SD = 3.62 | $t(18) = 6.12, p < 0.001$ |
| No reason vs Active separation | M = 59.31, SD = 3.47 | M = 56.52, SD = 5.29 | $t(18) = -0.68, p = 0.09$ |
| No reason vs Passive separation | M = 79.55, SD = 4.74 | M = 56.76, SD = 5.35 | $t(18) = 18.87, p < 0.001$ |

**Supplementary Table 4.** Classification performances for the contrasts between problems faced by the infants, for the model trained on infants’ problems and tested on maternal response, with mean (M) and standard deviation (SD) for both empirical and baseline distributions, and associated *t* statistics.

|  | Empirical | Baseline | Statistics |
| --- | --- | --- | --- |
| Pain vs Conflict | M = 71.00, SD = 13.29 | M = 55.20, SD = 2.99 | $t(78) = 7.34, p < 0.001$ |
| Social danger vs Passive separation | M = 70.71, SD = 14.22 | M = 55.17, SD = 3.14 | $t(78) = 6.75, p < 0.001$ |
| Social danger vs No reason | M = 64.32, SD = 14.90 | M = 54.50, SD = 3.17 | $t(78) = 4.08, p < 0.001$ |
| No reason vs Passive separation | M = 67.54, SD = 12.26 | M = 54.46, SD = 2.99 | $t(78) = 6.56, p < 0.001$ |
| Pain vs Active separation | M = 68.36, SD = 10.57 | M = 55.82, SD = 2.66 | $t(78) = 7.28, p < 0.001$ |
| Pain vs Passive separation | M = 70.38, SD = 10.26 | M = 55.03, SD = 2.99 | $t(78) = 9.09, p < 0.001$ |
| Social danger vs Active separation | M = 64.85, SD = 9.46 | M = 54.90, SD = 2.98 | $t(78) = 6.34, p < 0.001$ |
| Social danger vs Conflict | M = 61.54, SD = 10.47 | M = 55.29, SD = 2.90 | $t(78) = 3.64, p < 0.001$ |
| Pain vs No reason | M = 59.17, SD = 9.29 | M = 55.23, SD = 2.82 | $t(78) = 2.57, p < 0.01$ |
| Active separation vs Passive separation | M = 60.88, SD = 8.47 | M = 54.91, SD = 2.91 | $t(78) = 4.22, p < 0.001$ |
| Conflict vs Active separation | M = 61.17, SD = 5.40 | M = 54.28, SD = 2.59 | $t(78) = 7.27, p < 0.001$ |
| Conflict vs Passive separation | M = 57.44, SD = 7.00 | M = 55.47, SD = 2.91 | $t(78) = 2.91, p < 0.05$ |
| Pain vs Social danger | M = 61.25, SD = 5.91 | M = 54.51, SD = 2.91 | $t(78) = 6.47, p < 0.001$ |
| No reason vs Active separation | M = 56.69, SD = 5.91 | M = 55.51, SD = 3.17 | $t(78) = 1.31, p = 0.1$ |
| Conflict vs No reason | M = 55.07, SD = 3.95 | M = 55.24, SD = 2.66 | $t(78) = -0.23, p = 0.59$ |

**Supplementary Figure 1.**
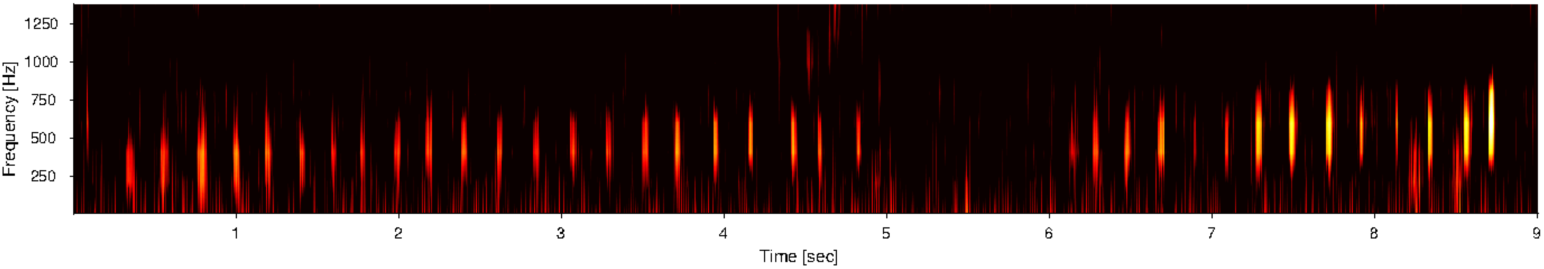
Time-frequency spectrogram of a series of distress calls given by individual OZ. The x-axis represents time, in seconds; the y-axis represents the frequency, in hertz.

**Supplementary Figure 2.**
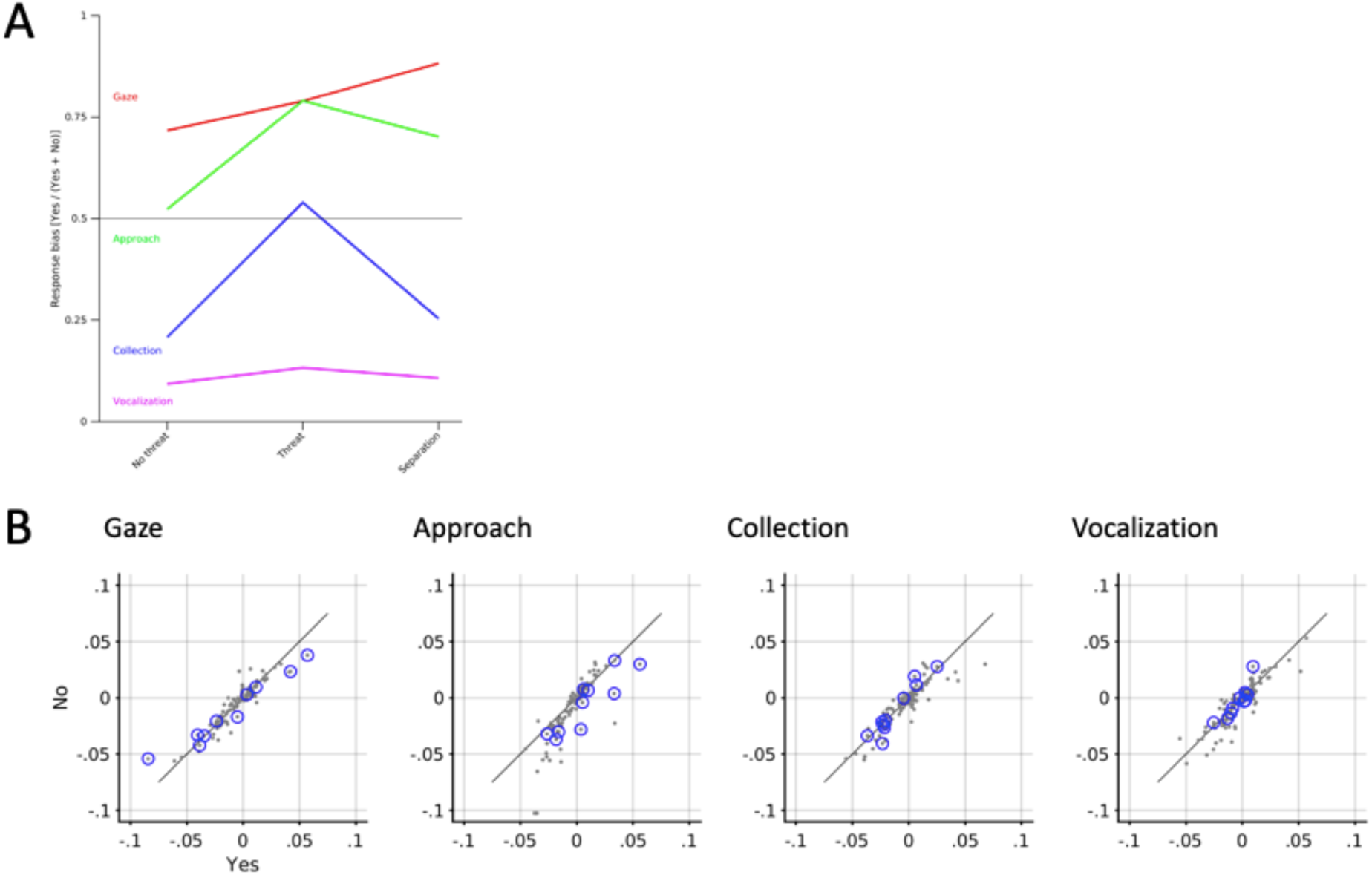
(A) Proportion of maternal responses across situations (B) Feature evaluation procedure, showing the percentage of feature dimensions with *p*-values smaller than .05 for each type of maternal response. Blue circles indicate selected features.

**Supplementary Figure 3.**
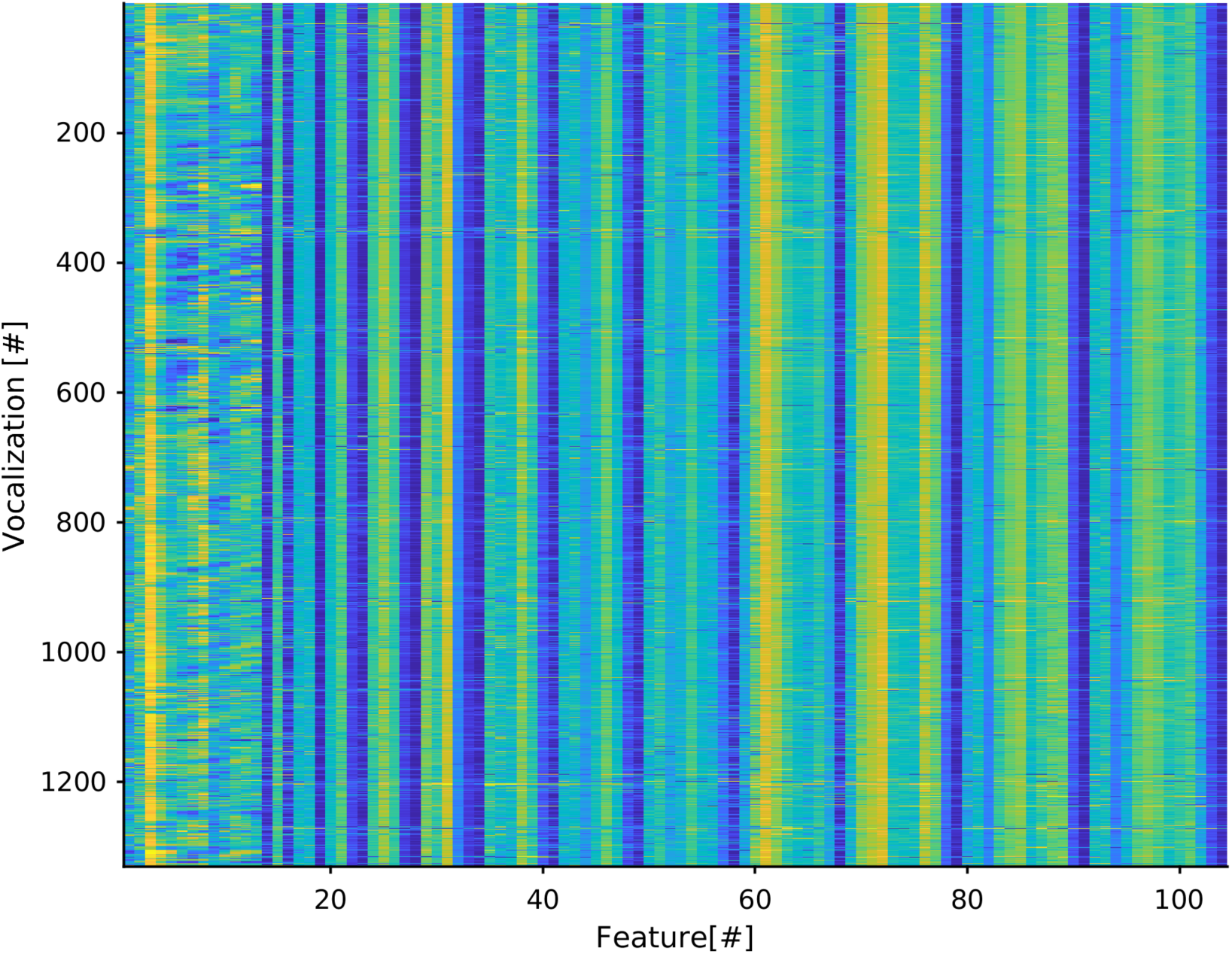
Feature extraction. Feature dimensions are shown in a colour representation: higher values are red, lower values blue. The first 13 values on the x-axis show the mean cepstra, the remain values the co-variances of cepstra.

